# Investigating the mutations in the SARS-CoV-2 proteins among European countries

**DOI:** 10.1101/2022.06.23.497239

**Authors:** Mohammad Abavisani, Karim Rahimian, Reza khayami, Mansoor Kodori, Mahsa Mollapour Sisakht, Mohammadamin Mahmanzar, Zahra Meshkat

**Affiliations:** Department of Microbiology, School of Medicine, Mashhad University of Medical Sciences, Mashhad, Iran; Bioinformatics and Computational Omics Lab (BioCOOL), Department of Biophysics, Faculty of Biological Sciences, Tarbiat Modares University, Tehran, Iran; Department of Medical Genetics and Molecular Medicine, Faculty of Medicine, Mashhad University of Medical Sciences, Mashhad, Iran; Non communicable Diseases Research Center, Bam University of Medical sciences, Bam, Iran; Department of Biochemistry, Erasmus University Medical Center, P.O. Box 2040, 3000 CA Rotterdam, The Netherlands; Department of Bioinformatics, Kish International Campus University of Tehran, Kish, Iran; Department of Microbiology and Virology, School of Medicine, Mashhad University of Medical Sciences, Mashhad, Iran

## Abstract

Severe acute respiratory syndrome coronavirus 2 (SARS-CoV-2) is a new member of the Coronaviridae family, triggering more than 190 million cases and more than two million deaths in European societies. Emerging the new variants due to mutations in genomic regions is foremost responsible for influencing the infectivity and mortality potential of such a virus. In the current study, we considered mutations among spike (S), envelope (E), membrane (M), and nucleocapsid (N) proteins of SARS-CoV-2 in the Europe continent by exploring the frequencies of mutations and the timeline of emerging them. For this purpose, Amino-acid sequences (AASs) were gathered from the GISAID database, and Mutation tracking was performed by detecting any difference between samples and a reference sequence; Wuhan-2019. In the next step, we compared the achieved results with worldwide sequences. 8.6%, 63.6%, 24.7%, and 1.7% of S, E, M, and N samples did not demonstrate any mutation among European countries. Also, the regions of 508 to 635 AA, 7 to 14 AA, 66 to 88 AA, and 164 to 205 AA in S, E, M, and N samples contained the most mutations relative to the total AASs in both Europe AASs and worldwide samples. D614G, A222V, S477N, and L18F were the first to fifth frequent mutations in S AASs among European samples, and T9I, I82T, and R203M were the first frequent mutations among E, M, and S AASs of the Europe continent. Investigating the mutations among structural proteins of SARS-CoV-2 can improve the strength of therapeutic and diagnostic strategies to efficient combat the virus and even maybe efficient in predicting new emerging variants of concern.

## Introduction

After severe acute respiratory syndrome coronavirus (SARS-CoV) and Middle East respiratory syndrome–related coronavirus (MERS-CoV) outbreaks in 2003 and 2012, respectively, the SARS-CoV-2 is the third beta coronavirus responsible for an outbreak accounting for more than 190 million cases and more than two million deaths in Europe [1, 2]. SARS-CoV-2 has a 26-32kb positive-sense single-stranded RNA genome with 79.5% similarity to SARS-CoV [3, 4]. The SARS-CoV-2 genome has 14 open reading frames (ORFs) encoding various proteins, the function of which is not fully understood. The proteins are categorized into three groups: structural, non-structural, and accessory proteins [5]. The 5’ tail of the genome is where pp1a/ab ORF exists cleaved to 16 non-structural proteins [5]. The 3’ part of the genome encodes the structural proteins, including the spike glycoprotein (S), the envelope protein (E), the membrane protein (M), and the nucleocapsid (N). The last part of the genome encodes the accessory proteins (3a, 3b, p6, 7a, 7b, 8b, 9b, and orf14) [6]. While the non-structural proteins are mainly involved in replicating the virus, the structural proteins are essential for pathogenicity and immune response evasion. For example, the S protein helps the virus enter the host cells via angiotensin-converting enzyme 2 (ACE2) [7]. Therefore, changes in these proteins could vastly alter the virus characteristics.

RNA virus replication has a high error rate. This could be considered a double-edged sword for the virus as it could result in the adaptation of the virus to different environments or its extinction dependent on the location and types of mutations [8]. Although Coronaviruses have proofreading ability, several studies have reported various mutations in the SARS-CoV-2 genome. These mutations have conferred more immune resistance and pathogenicity to the virus and have provided challenges in diagnosis, treatment, and prevention of the disease. For example, the D614G mutation in the S protein, first observed in China and then in Europe, has increased virus infectivity and transmissibility [9].

The emergence of mutations over time has resulted in five variants of concern (VOC), namely Alpha, Beta, Gamma, Delta, and Omicron, where mutations in the receptor-binding domain (RBD) of the S protein substantially increased its binding affinity and infectivity [10]. Recognizing mutations’ evolutionary patterns could help identify novel therapeutic or vaccination approaches or even predict a new VOC. Thus, our study analyzed the evolutionary trends of SARS-CoV-2 mutations in Europe and identified the frequencies of mutations within structural proteins between January 2020 and April 2022.

## Materials and Methods

### Sequence gathering

Amino-acid sequences (AASs) of structural proteins of SARS-CoV-2 from January 2020 until April 2022 were extracted from GISAID database (https://www.gisaid.org/) under the permission and monitoring of Erasmus Medical Center [11-13]. Wuhan-2019 virus with access number of EPI_ISL_402124 was utilized as reference sequence in comparison process. Comparison process was operated among S region with 1273 AA, AASs of E region with 75 AA, M region with 222 AA, and N region with 419 AA. The sequences being from non-human samples or own unspecified time or location were excluded from the study. Also samples with different AAs length compared to reference sequence and non-specified AAs did not involve in the current study.

### Sequences alignment and mutation tracking

Sequences alignment was conducted using Python 3.8.0 software by FASTA files. Mutation tracking was performed via detecting any difference between samples and Wuhan-2019. In the following, data optimization was accomplished using ‘Numpy’ and ‘Pandas’ libraries. The algorithm applied to mutant finding is written in the supplementary file 1. Due to the identical lengths among all sequences by the virtue of excluding samples with divergent size, the mentioned algorithm used ‘Refseq,’ and ‘seq’ refer to Wuhan-2019 sequence and sample sequence, individually. Final reports include mutations with attributed locations and subset AA.

### Data normalization and statistical analysis

By the virtue of R 4.0.3 and Microsoft Power BI, data normalization was conducted to achieve the high quality comparison. For that reason, the number of mutations was divided by the number of attributed sequences that were comparable in equal proportions for each of the European countries.

## Results

### Prevalence and incidence rate of mutations

13184933 European samples were imported to the study included 328188 samples for S AASs, 5134947 for E AASs, 4601354 for M AASs and 3120444 samples for N AASs. Besides, the numbers of AASs qualified globally to considering in the current study were as 950459, 9914529, 8860463 and 6365457 samples belonging to S, E, M and N AASs, respectively.

Studying the frequency of mutations among structural proteins among European countries demonstrated that 8.6% of S samples did not carry any mutation and 19.6%, 34.2%, 21.7% and 15.8% of them displayed one, two, three and more than three mutations, respectively (Figure 1A). In comparison, 4.8% of worldwide AASs demonstrated no mutation and 26.3%, 25.3%, 13.7% and 29.8% contained one, two, three and more than three mutations, respectively. On the other hand, 63.6%, 24.7% and 1.7% of E, M and N samples did not demonstrate any mutation in Europe continent, respectively (Figure 1B-D). In this continent, 36.2% of E AASs carried one mutation and 0.1% of them displayed two mutations. Also 44.8% of M samples in Europe showed one mutation and 14.9% showed two mutations. The frequency rates of one and two mutations among N AASs from Europe were inferred as 5.2% and 4.5%, respectively.

**Figure 1.**
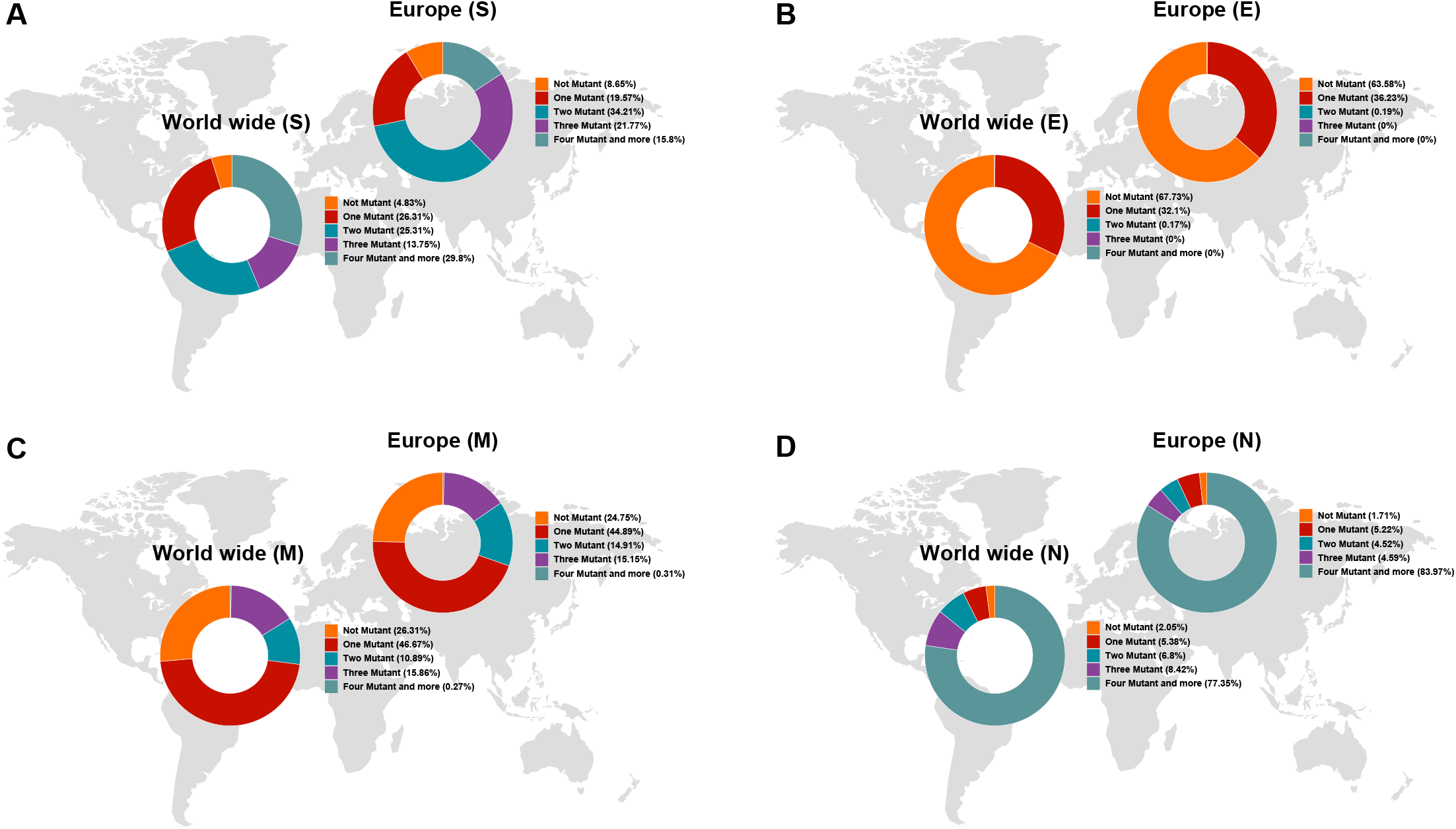
Pie chart plot of the number of mutations of S, E, M, and N proteins in SARS-COV-2 as of April 2022 in Europe, and Worldwide. Sections A, B, C, and D display data belonging to S, E, M, and N proteins, respectively.

In order to detecting the regions of AASs with high mutation predisposition, the heat map was designed. According to the heat map, the most mutations relative to the total AASs among the S, E, M and N AASs from both European samples and global data were occurred in the regions of 508 to 635 AA (0.0074 frequency in Europe and 0.0077 frequency globally), 7 to 14 AA (0.0501 frequency in Europe and 0.0438 frequency globally), 66 to 88 AA (0.0208 frequency in Europe and 0.0219 frequency globally) and 164 to 205 AA (0.0295 frequency in Europe and 0.0292 frequency globally), respectively (Figure 2). In consideration the mutation frequencies among regions of S AASs in European samples, the regions of 127 to 254 (0.0039 frequency) and 1 to 127 (0.0027 frequency) were concluded as the second and third region. Second and third regions in heat map of other structural AASs among European samples were respectively inferred as 56 to 63 (0.0004 frequency) and 49 to 56 (0.0002 frequency) regions for E AASs, 1 to 22 (0.0203 frequency) and 44 to 66 (0.0131 frequency) regions for M AASs and 205 to 246 (0.0226 frequency) and 41 to 82 (0.0154 frequency) regions for N AASs.

**Figure 2.**
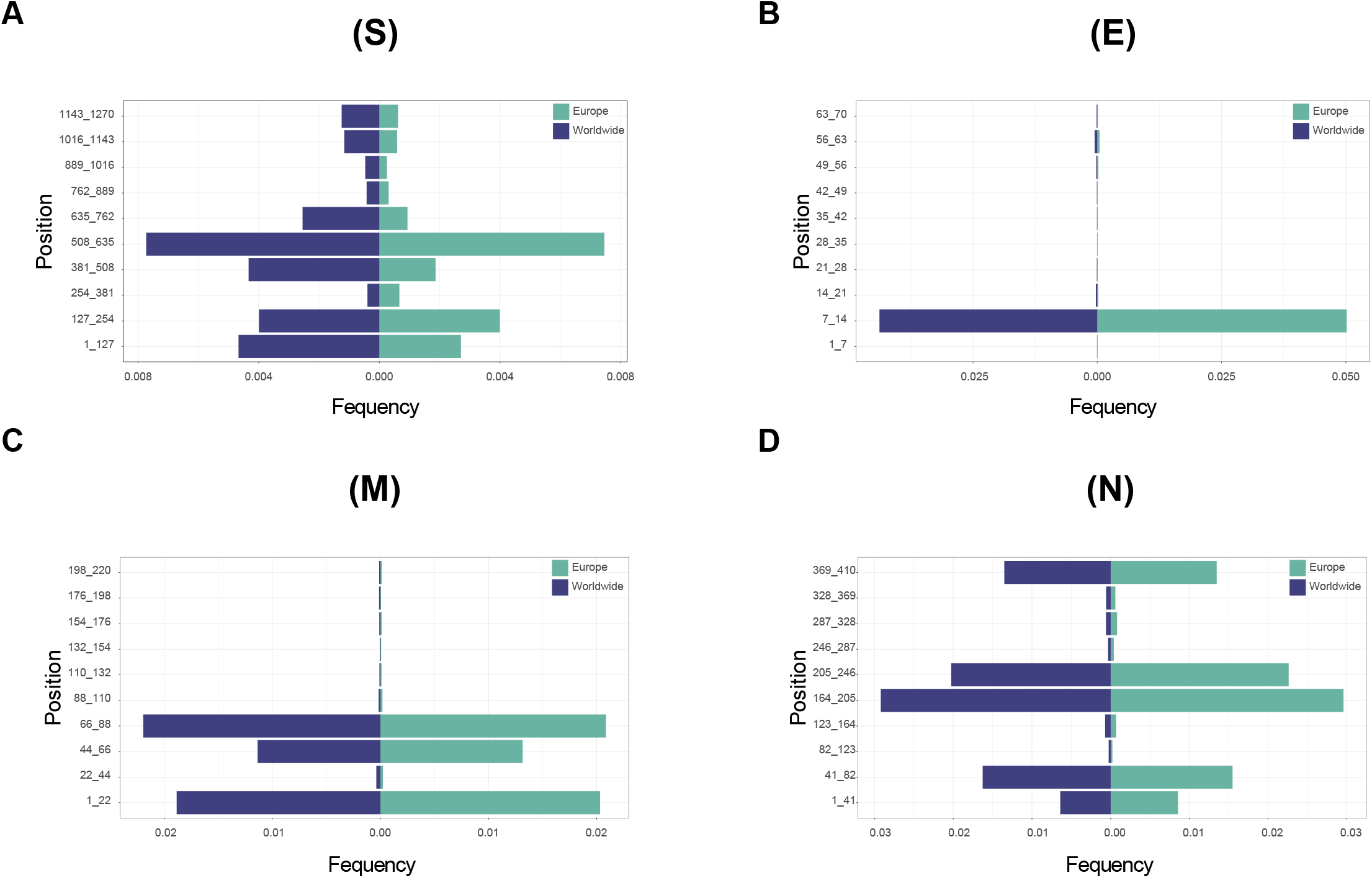
Hotspot regions in S, E, M, and N proteins of SARS-COV-2 as of April 2022. The plots indicate the rate of mutation per 100 AAs in Europe and Worldwide. The hotspot regions of S, E, M, and S were displayed in the sections of A, B, C, and D, respectively.

### Structural substitutions and frequencies

To understand the location of occurred mutation among structure of protein was another investigating step in the current study. So, data including locations beside their frequencies were gathered from the beginning of the pandemic to April 2022. D614G with 0.9756 frequency rate was concluded as the most frequent mutation among S AASs of European samples (Figure 3A). In the following, A222V (0.3895 frequency rate), S477N (0.1600 frequency rate), L18F (0.1575 frequency rate) and E488K (0.0304 frequency rate) were second to fifth frequent mutations in S AASs, respectively. Moreover, N501Y with 0.0974 frequency rate, H655Y with 0.0957 frequency rate, P681H with 0.0852 frequency rate, K417N with 0.0680 frequency rate and T478K with 0.0761 frequency rate were observed as the second five frequent mutations among S AASs, respectively. Data inserted from worldwide samples displayed D614G (0.9764 frequency), E484K (0.1406 frequency) and L18F (0.1623 frequency), A222V (0.1496 frequency) and N501Y (0.1355 frequency) as first to fifth frequent mutations among S AASs. On the other hand, T9I (0.3508 frequency rate), P71L (0.0035 frequency rate) and V58F (0.0013 frequency rate) were inferred as the first three frequent mutations in E AASs from European samples, respectively (Figure 3B). Despite the first and the second frequent mutations in E AASs of global data were identical with European samples, they displayed different frequencies. The worldwide samples introduced T9I (0.3064 frequency), P71L (0.0046 frequency) and V62F (0.0022 frequency) as first to third frequent mutations in E AASs. Among European samples, all of the fourth to eighth frequent mutations in E AASs demonstrated substituting AA to phenylalanine (F). These were resulted as V62F with 0.0012 frequency rate, S55F with 0.0009 frequency rate, L21F with 0.0009 frequency rate, L73F with 0.0008 frequency rate, and S68F with 0.0006 frequency rate, respectively. Moreover, V24M with 0.0002 frequency rate and A41V with 0.0002 frequency rate were displayed as ninth and tenth frequent mutations, respectively.

**Figure 3.**
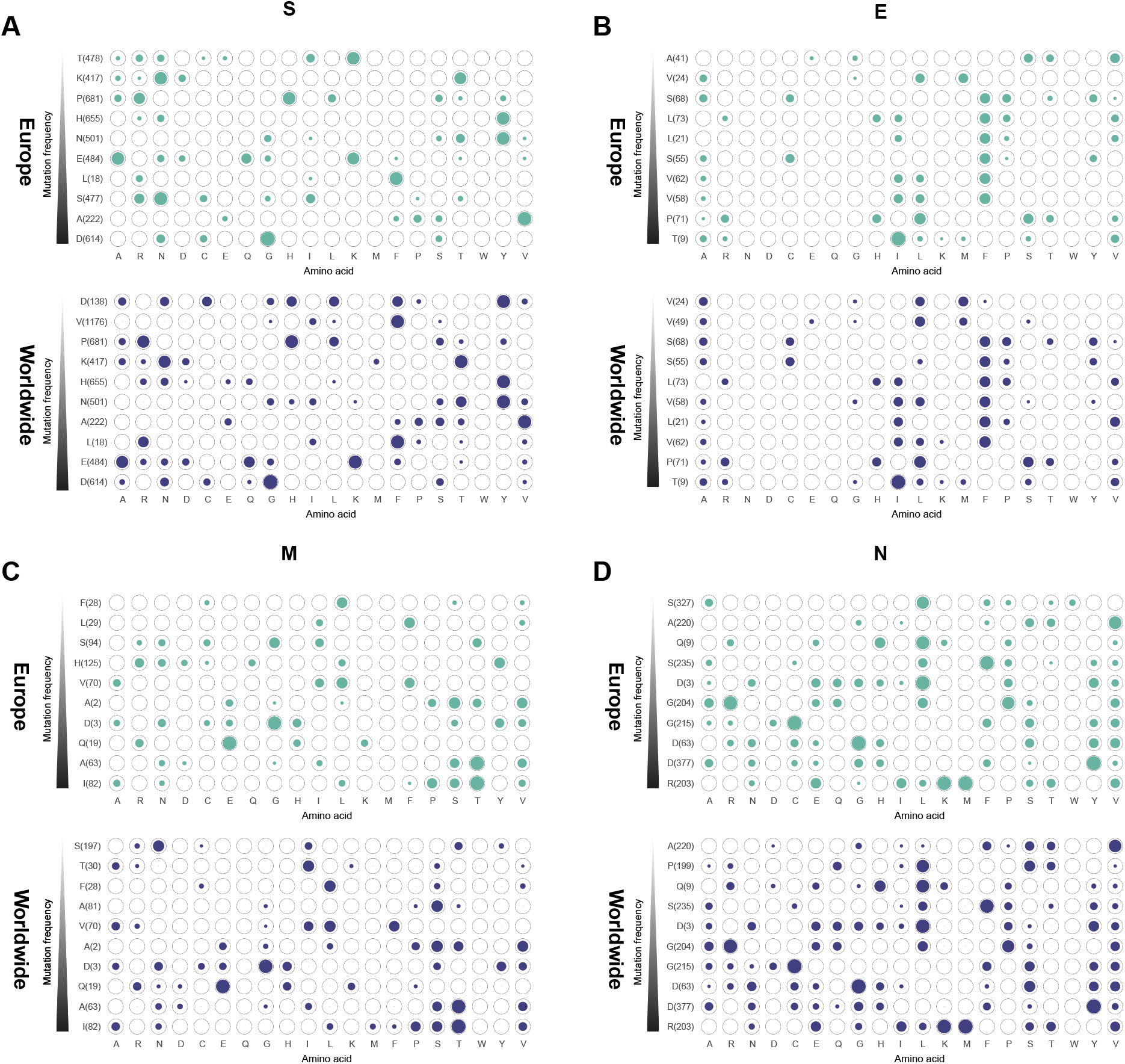
Top 10 mutations in S, E, M, and N proteins of SARS-COV-2 with the highest frequency in Europe and Worldwide. The position of altered AASs and substituted ones is shown differently based on the frequency percentage of substituted AA. The mutation frequency was estimated for each of them by normalizing the number of genomes carrying a given mutation in a desired geographic area.

Additionally, data gathered from M AASs among European samples displayed that I82T (0.4504 frequency rate), A63T (0.2878 frequency rate), Q19E (0.2860 frequency rate), D3G (0.1533 frequency rate) and A2S (0.0038frequency rate) were first to fifth frequent mutations, respectively (Figure 3C). Also results demonstrated that V70L, H125Y, S94G, L29F and F28L were sixth to tenth frequent mutations in M AASs with 0.0020, 0.0012, 0.0010, 0.0010 and 0.0010 frequency rates, respectively. Data achieved globally showed identical arrangements of first to fourth frequent mutations with European samples among M AASs in spite of different frequencies. Consideration data of N AASs inferred that the first frequent mutation was observed in the location of R203, which displayed as R203M/K mutations with 0.6298 / 0.2674 frequency rates (Figure 3D). Furthermore, D377Y (0.6326 frequency rate), D63G (0.6226 frequency rate), G215C (0.5963 frequency rate) and G204R (0.2425 frequency rate) were second to fifth frequent mutations in N AASs. Besides, results exhibited second five frequent mutations among N AASs as D3L (0.2344 frequency rate), S235F (0.2348 frequency rate), Q9L (0.0825 frequency rate), A220V (0.0471 frequency rate) and S327L (0.0192 frequency rate), from sixth to tenth respectively. Additional data about the frequencies of mutations in European samples and worldwide AASs were listed in supplementary files Frequencies-Europe.xlsx and Frequencies-Worlwide.xlsx.

### Evolutionary trends of top ten mutations based on time

Analyzing the evolutionary trends of mutations among structural proteins of SARS-CoV-2 detected that mutation among aspartic acid (D) in the location of 614 among European and total S AASs S was occurred from January 2020 but gained to decrease and became increased again from February 2020(Figure 4A). From August 2020 till April 2022, the frequencies of D614G mutations were almost identical and in a steady trend. In comparison, August 2021 was observed as a time that frequencies of some mutations among S AASs became changed. For instance, E484K and N501Y mutation gained to decrease. These trends were in contrast to P681R and T478K mutations which gained to increase from August 2021. Conversely, timeline data belonging to E AASs demonstrated similar trends of distribution for structural mutations excluding T9I mutation (Figure 4B). Steady trend of T9I mutation was changed and gained to increase from November 2021 and has been received to maximum frequency after February 2022.

**Figure 4.**
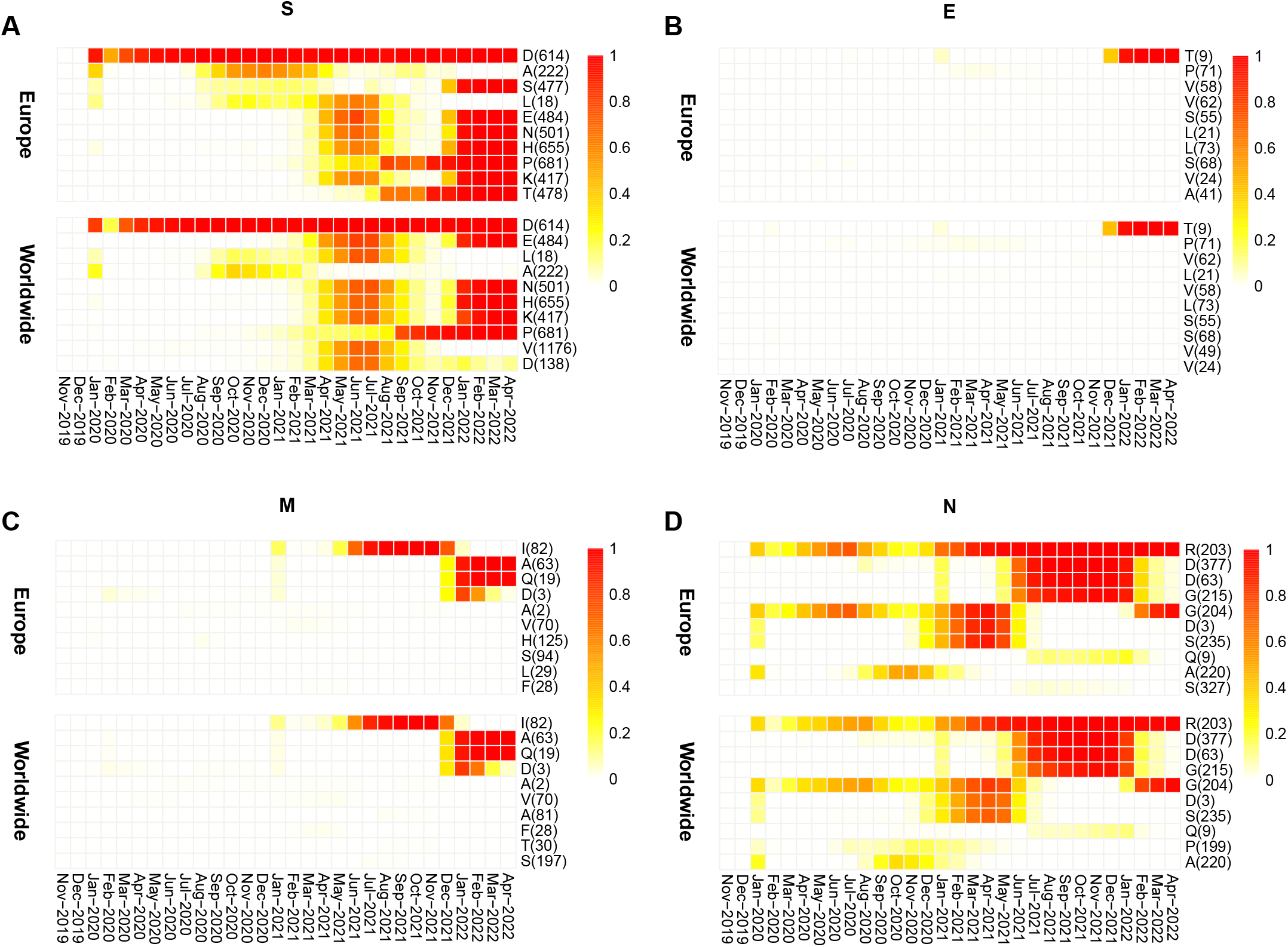
Time evolution heat maps of top ten high-rate mutations of S, E, M, and N proteins of SARS-COV-2 in different geographic areas including Europe and Worldwide. According to the month of sample collection, data is computed as the number of AASs having a given mutation over the total number of AASs according to the month of sample collection.

Frequency trends of I82T mutation, which had the most frequent mutation among M AASs, were in steady and changeless form till January 2021 (Figure 4C). In contrast, distribution of I82T mutation gained to increase after this time and after meeting with its own highest frequency in September to November 2021, the frequency is noticeably reduced. Data displayed that after the time when I82T frequency gained to decrease, the frequencies of A63T and D3G gained to increase. The differences between these two latter mutations was observed in their increasing trends, in which D3G frequency gained to decrease from January 2022 but A63T sustained its own increasing trend of mutation till the time of study. The frequency of mutations among N AASs was concluded highly fluctuant (Figure 4D). In an overview, it seems arginine mutation (R) in the location of 203 had a growing evolutionary trend in spite of oscillating frequencies. Additionally, frequency trends of D377Y, D63G and G215C had similar pattern with increasing tendency from February 2021 and decreasing movement from 9 months later, November 2021. Further details of timeline data were displayed into the supplementary files named Timeline-Europe.xlsx and Timeline-Worldwide.xlsx.

## Discussion

As the SARS-CoV-2 virus evolves and evolves into COVID-19 in this ongoing pandemic, monitoring its evolution is critical to developing new diagnostic tests, vaccines, and treatments. When analyzing the number of mutations, we found that the S and the N proteins have undergone the most alterations. We found that the distribution of mutations in the S protein differs between Europe and the rest of the world. The majority of samples in Europe had only two mutations (34.21%) in the S protein, while in the worldwide samples, the majority of the samples had four mutations and more (29.8%). The majority of samples in both Europe and worldwide groups had four mutations and more in the N protein. On the other hand, the distribution of mutations in the E and the M protein was the same in Europe and Worldwide. In the E protein, more than 60% of the samples did not have any mutations, and more than 30% had only one mutation, T9I, which has emerged in the Omicron variant. Finally, more than 70% of the samples had at least one mutation in the M protein.

The SARS-CoV-2 genome has been evolving, and the S protein of the virus is dominant in mutations. This protein is the main contributor to cell binding and entry, thus impacting viral infectiousness [14]. Interestingly, The D614G variation in the S protein of SARS-CoV-2 is the most frequent mutation in the viral genome. Korber et al. showed that D614G is associated with lower RT-PCR cycle thresholds suggesting an increase in the viral load of SARS-CoV-2 in the upper respiratory tract [15]. They further indicated that D614G could disrupt a hydrogen bond formed between the aspartic acid in residue 614 of the S1 subunit and a threonine located at position 859 of the S2 subunit of the S protein and possibly affect glycosylation in the adjacent N616 site [15]. This phenomenon is suggested to make the S protein more stable and facilitate the S protein incorporation into the pseudovirion [16]. Consistent with this, Li et al. showed that pseudotyped viruses with D614G had increased viral infectivity [17].

Furthermore, we found that the prevalence of the two of the most prevalent mutations found in the S protein (A222V, L18F) in Europe and the world was decreased recently, possibly due to decreased advantage in the process of viral evolution. On the other hand, the frequency of K417N S477N, T478K, E484A, N501Y, H655Y, and P681H is increased recently, among which S477N and T478K were found only in Europe’s top ten mutations. K417N S477N, T478K E484A, and N501Y are located in the spike protein’s receptor-binding domain (RBD) and may heavily impact the SARS-CoV-2 host cell entrance ability [18]. In fact, it has been shown that change of Asparagine (N) to Tyrosine (Y) in the position 501 of the S protein results in increased binding affinity to ACE2 receptor, increasing transmission ability [18]. K417N and E484A are significantly disruptive, making Omicron more likely to have vaccine breakthrough [19]. S477N was the primary mutation in the spike protein of 20A.EU2 in fall 2020 in Europe and has been found to resist neutralization by monoclonal antibodies (mAb) and convalescent sera [20, 21]. According to Chen et al., S477N and T478K are dependent on each other [19]. Although the information on the physiological impact of T478K is conflicting, it is clear that the variant enhances infectivity [22]. Also, it could be effective in helping the virus evade antibody neutralization and impairing the interaction of RBD with drugs [23, 24]. We found that the Glutamic acid located in position 484 of the S protein often changes to A, K, and Q in both Europe and the World. Liu et al. studied the effect of 19 mAbs against the RBD of S protein and found that E484K could escape neutralization by convalescent sera [25]. E484A was seen in the Omicron variant and exerts a similar immune evading effect as E484K seen in the Beta variant [26]. Also, it has been found that E484Q alongside L452R could synergistically facilitate immune evasion in B.1.617.1 and B.1.617.3 [27].

Furthermore, N501Y has been shown to enhance viral transmission compared to the wild-type [28, 29]. One reason could be the fact that this mutation could increase the binding affinity of RBD-ACE2 by ten folds facilitating cell host entry [30, 31]. Two other critical mutations are H655Y and P681H, which are not located in the RBD and enhance the S protein cleavage possibly contributing to viral transmission [32, 33]. These mutations have substantial effects on vaccine efficacy and immune evasion. For example the variants containing E484K, K417N, and N501Y mutations have been shown 25%-50% increase in evading vaccine immunity [34].

Other structural proteins of SARS-CoV-2 have been less studied, possibly because they contain fewer effective mutations in their sequences. Nevertheless, studying these structural proteins may help develop more effective therapeutic and diagnostic measures. We found that the E protein has been more conserved in the evolution process of SARS-CoV-2, with only T9I mutation as the significant change in this protein. Consistently, other studies have reported this mutation as the most prevalent mutation in the E protein of the Omicron strain [35, 36]. Although its function is not entirely understood, the change in the 9^th^ position of E protein has been associated with stronger interaction of the virion with membrane lipids [37].

On the other hand, the M protein has been subject to more changes. The I82T mutation has been the most frequent change in the M protein, although, since December 2021, its frequency has decreased, and other mutations such as A63T and Q19T have shown increased prevalence recently, possibly because they confer a fitness advantage to the virus. We found that substitution of Arginine (R) to Lysine (K) or M in the 203rd position of the N protein is the most frequent mutation in this protein in Europe and the world. Interestingly, this mutation has been observed in Alpha, Delta, and Omicron lineages [38, 39], confers resistance against immune response, and increases infectivity [38-41]. This mutation facilitates virion assembly by increasing the condensation of N-protein with viral genomic RNA, hence gaining more frequency [42]. Another interesting mutation is G204R in Europe and the rest of the world. This substitution was one of the most common mutations alongside R203 till May 2021; however, its prevalence decreased from May to November 2021, and with the emergence of the Omicron variant, its frequency increased again. Apparently, variants containing 203K/204R have enhanced ribonucleocapsid assembly ability and are more infectious than the wild-type variant [43].

All in all, our findings provide valuable information to understand evolutional trends of SARS-CoV-2 virus and contribute to the control of the pandemic and developing new strategies such as new vaccine targets and drugs. Our study was limited in that we focused on the AAS of the viral genome overlooking the nucleotide sequence and codon bias. Further studies are required to address the functional effects of these structural mutations and monitor the evolutional trends.

## Supporting information

Frequencies-Europe.xlsx

Frequencies-Worlwide.xlsx

Timeline-Europe.xlsx

Timeline-Worldwide.xlsx

## Acknowledgments

The authors thank all of the researchers who have shared genome data openly via the Global Initiative on Sharing All Influenza Data (GISAID).

## Additional information

### Supplementary Information

Supplemental data associated with the current study have been gathered.

